# The Transcription Factor Sob Maintains Intestinal Stem Cell Homeostasis to Delay Ageing

**DOI:** 10.64898/2026.05.29.728682

**Authors:** Fanila Shahzad, Wenbin Wei, Anna L. M. Smith, Laura C. Greaves, David P. Doupé

## Abstract

Ageing is associated with physiological decline and disrupted tissue homeostasis, often driven by imbalanced stem cell activity. The *Drosophila* midgut serves as a powerful model for studying epithelial homeostasis and ageing as it exhibits many conserved hallmarks of regulation and deteriorating intestinal stem cell function with age. We have identified the transcription factor encoded by *Sister of odd and bowl* (Sob) as a conserved marker of intestinal stem/progenitor cells in homeostasis whose expression is lost with age. Premature downregulation of *sob* shortens lifespan while overexpression in the intestinal stem/progenitor cells is sufficient to extend lifespan, suggesting a critical role in maintaining homeostasis with age. At the cell and tissue level stem/progenitor knockdown and overexpression reveal that Sob maintains homeostasis by regulating both proliferation and lineage specification. In young guts, Sob restricts intestinal stem cell proliferation and promotes enteroendocrine differentiation while limiting excessive enterocyte differentiation. We have also found that the transcription factor Exex, the Notch pathway inhibitor *Numb*, and the enterocyte differentiation regulator Dawdle act downstream of Sob to regulate proliferation and differentiation. Our findings position Sob as a critical molecular switch linking stem cell dysfunction to gut ageing.

## Introduction

The mammalian intestinal epithelium is rapidly turned over throughout adult life. Intestinal stem cell (ISC) proliferation replaces the continuous loss of differentiated cells into the lumen, and is tightly regulated to maintain homeostasis (reviewed^1^). Similar processes occur in the *Drosophila* midgut epithelium, with ISCs replacing absorptive enterocytes and secretory enteroendocrine cells during normal homeostasis^2,3^. Many of the signalling pathways and regulatory mechanisms are conserved from flies to mammals making them an excellent model system^4,5^. In the ageing gut of both mammals and flies this homeostatic balance breaks down. Stem cells undergo mis-regulated proliferation, loss of balanced differentiation and exhaustion leading first to dysplasia and ultimately a loss of tissue integrity^6-8^.

Intestinal ageing is a complex, multifactorial condition with disruption to signalling in the stem cell regulatory niche, local and systemic inflammation, dysbiosis of the microbiota and loss of barrier function all contributing and interacting^6^. Ultimately, at the stem cell level, these inputs converge on transcriptional regulation with the effector transcription factors of regulatory pathways such as EGFR, JAK/Stat, NF-Kß and JNK intersecting with cell type specifc transcription factors^6^. Conserved transcription factors including Escargot, Sox21a, Sox100B and the Enhancer of Split Complex (E(Spl)-C) have been shown to regulate normal stem cell proliferation and differentiation^9-17^. Escargot is widely used as a marker of ISCs and their enteroblast (EB) progeny and is important for their maintenance and differentiation^9,10^ while its ortholog SNAI1 is involved in mouse intestinal stem cell differentiation^l8^. Sox21a and Sox100B are required for EB to enterocyte (EC) differentiation and coordinate differentiation and proliferation^11-16^. Sox21a ortholog SOX9 regulates proliferation and Paneth cell differentiation^19,20^. The Enhancer of split complex genes are key Notch targets and promote differentiation, and ortholog ASCL2 promotes stem cell identity, possibly reflecting Notch’s opposing functions in *Drosophila* and mouse ISCs^17,21^. Conserved transcriptional regulation also extends to chromatin regulators with Chronophage, the *Drosophila* ortholog of SWI/SNF component BCL11B, required for ISC proliferation^22,23^.

Changes in expression of some transcriptional regulators including Caudal and Chronophage have been shown to influence ISC ageing^22,24^. However, the changes in transcriptional regulation of stem cell fate and how this intersects with niche signalling pathways in ageing are not fully understood. An improved mechanistic understanding of how homeostasis breaks down with age has the potential to identify novel biomarkers of ageing and possible interventions to slow the ageing process. Here we identify OSR2/Sob as conserved transcription factors marking intestinal stem and progenitor cells whose expression is reduced with age. Manipulation of Sob levels in *Drosophila* ISCs and their enteroblast progenitors impacts stem cell proliferation and differentiation, and organism lifespan. These effects are mediated through regulation of the Notch and TGF-ß pathways, and the transcription factor Exex. Our findings establish Sob as a key and conserved regulatory switch that is lost with age.

## Results

### Mammalian OSR2 and *Drosophila* ortholog Sob are expressed in ISCs and Progenitors

Analysis of previous expression profiling data found Sob, the ortholog of human OSR2^25^, is expressed in *Drosophila* intestinal epithelium and its levels change with age^26^. Human protein atlas data suggests that OSR2 is expressed in intestinal crypts making it a candidate marker of stem cells^27,28^. We therefore labelled mouse intestinal epithelium for OSR2 using immunohistochemistry and observed strong expression in the intestinal crypts consistent with expression in ISCs, Paneth cells and TA cells and also in secretory Goblet cells along the crypt villus axis (Figure 1A). To assess whether these expression patterns change with age we stained old and young mouse intestinal epithelium and saw a reduction in OSR2 expression in the crytps with age (Figure 1B). In parallel a *sob* GAL4 enhancer trap was used to drive expression of EGFP in *Drosophila* and showed specific expression in midgut cells with small nuclei co-expressing the ISC/EB marker esg-lacZ (Figure 1C). Use of additional markers to separately label ISCs (Delta antibody staining) and EBs (Notch transcriptional reporter, Su(H)GBE-lacZ) confirmed that sob is expressed in both stem and progenitor cells (Supp Figure S1A-E). Analysis in ageing flies showed a marked reduction in sob expression with age with around 80% of aged guts showing no expression of the sob-GAL4 reporter (Figure 1D, Supp Figure S1F). Together these results suggest that OSR2/Sob transcription factors may be conserved markers of homeostatic stem and progenitor/ transit-amplifying (TA) cells whose expression is lost with age.

**Figure 1:**
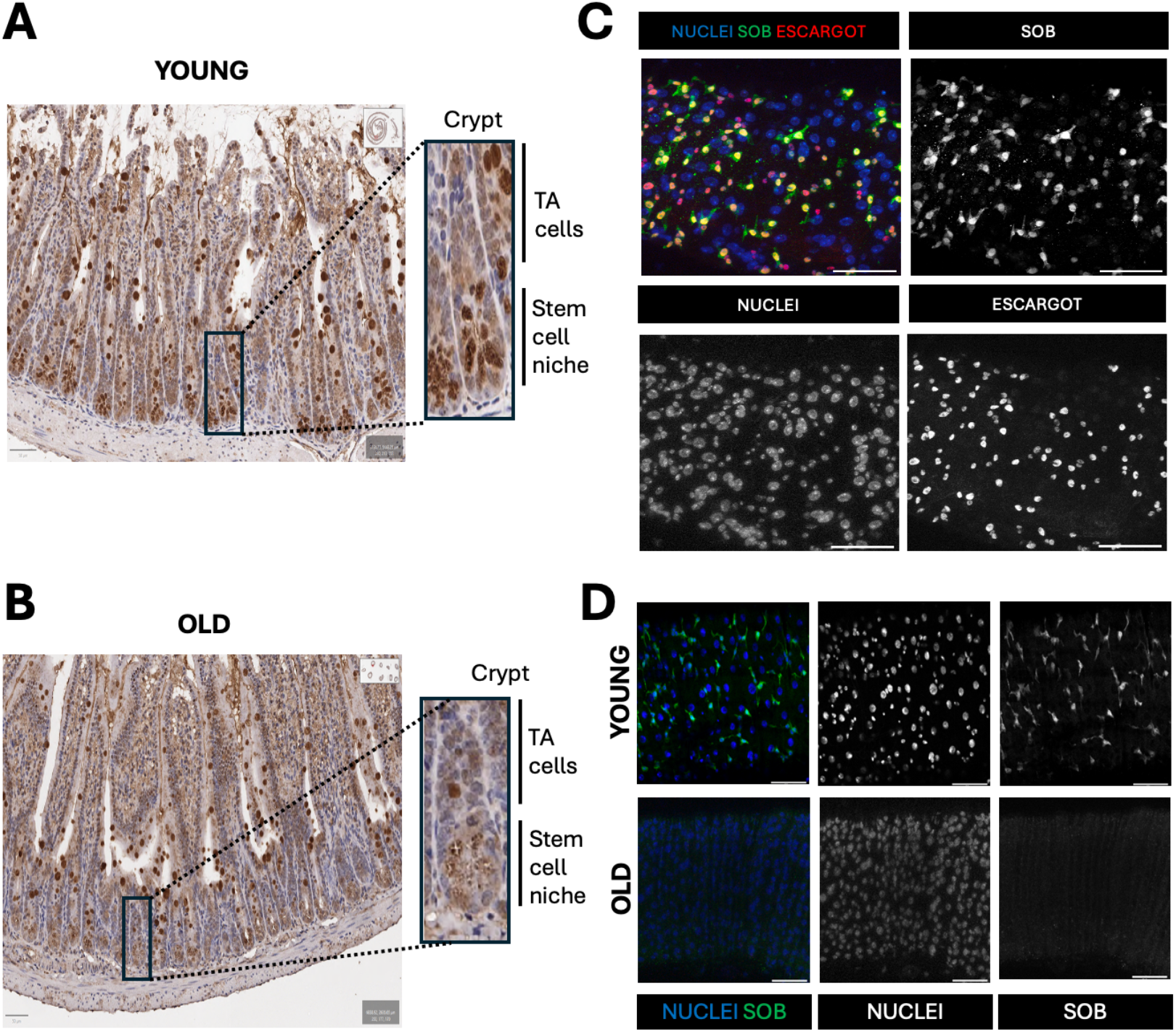
OSR2/Sob expression in Intestinal Homeostasis and Ageing. (A,B) Immunohistochemistry staining of young (13 weeks) (A) and old (16-18 months) (B) mouse small intestine. Zoomed images show individual crypts. n≥5 /condition (C, D) Representative maximum intensity-projected confocal z stack images of posterior midgut with brightness and contrast uniformly enhanced for clarity. (C) 10-day old sob-Gal4,UASeGFP with esgLacZ. Grey-scale images show nuclei (DAPI), sob expressing cells (sob-GAL4, UASeGFP), and ISC/EBs (esgLacZ). Merged images show nuclei (blue), sob (green), and ISC/EBs (red). (D) Young (10 days) and old (30 days) midgut showing nuclei (blue) and sob (green). Scale bars 50 µm.

### Intestinal Stem / Progenitor Sob regulates *Drosophila* lifespan and Intestinal Homeostasis

The timing of loss of Sob expression correlates with age-related decline in intestinal homeostasis and stem cell mis-regulation. We therefore assessed the effects of sob knockdown throughout adult life on lifespan using a drug-inducible, ISC/EB specific, geneswitch-GAL4 driver line, 5961^GS^. This enables knockdown and overexpression studies in a consistent genetic background at 25°C^29^. ISC/EB specific knockdown of *sob* led to a reduction in median lifespan from 62 days to 55 days (Figure 2A and Supplementary Figure S2). In the reciprocal experiment, sustained overexpression of Sob in ISC/EBs extended median lifespan from 50 to 57 day (Figure 2B and Supplementary Figure S2). A key feature of ageing at the stem cell level is misregulated proliferation with age. In control conditions a significant increase in mitotic index is observed by 30-40 days (Figure 2C). Overexpression of Sob has no effect on the relatively low midgut mitotic index in young flies but suppressed the age-related increase at 30 and 40 days (Figure 2C). This is consistent with Sob’s effect on lifespan being mediated by preserving stem cell proliferative homeostasis.

**Figure 2:**
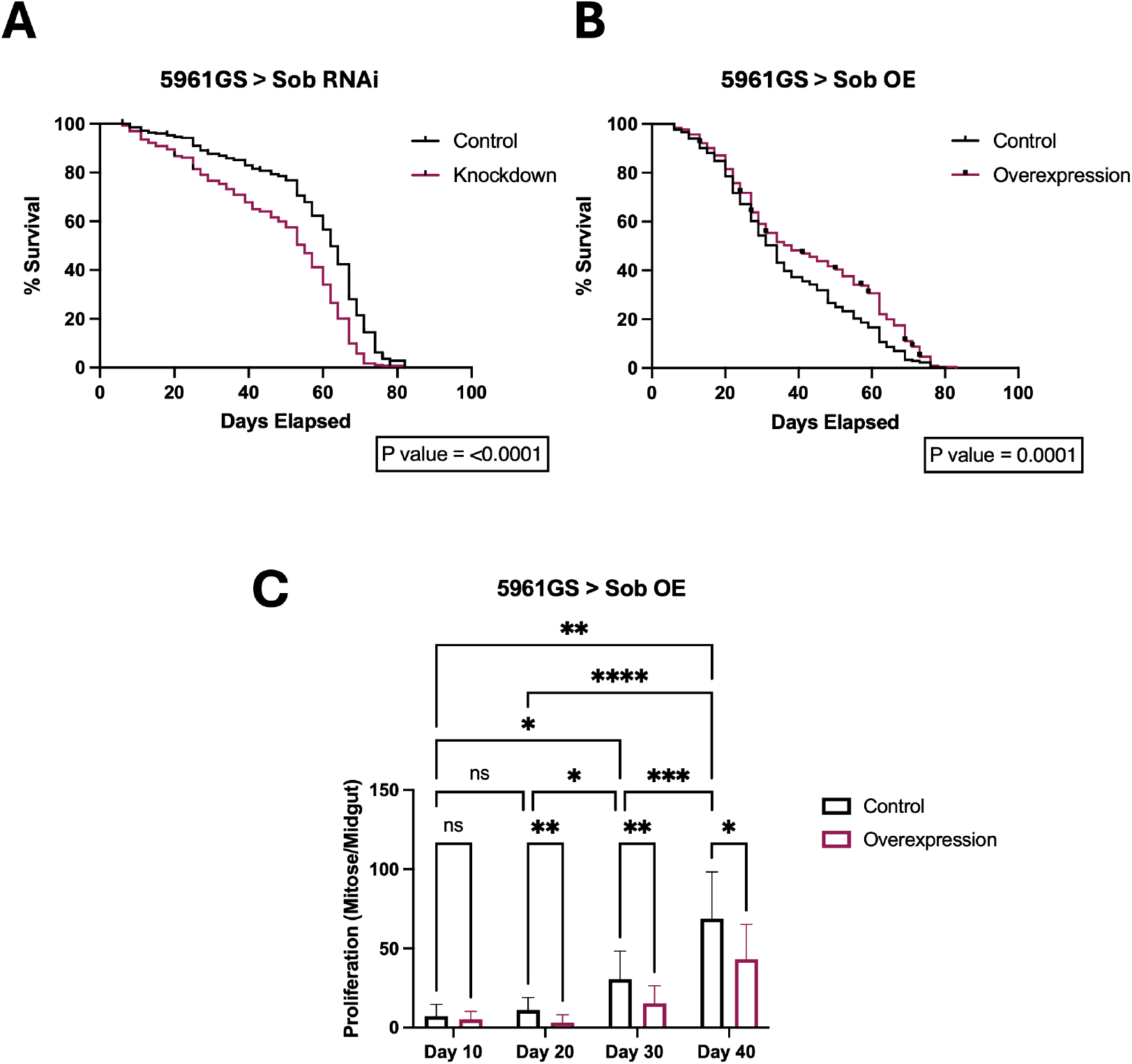
Stem/ progenitor Sob Impacts Lifespan. Lifespans of sob knockdown (A) and overexpression (B) using the driver 5961GS. p values (log-rank Mantel-Cox test) are indicated on each plot, n=300 flies/condition. (C) Effect of sob overexpression from Day 7 onwards on midgut mitosis. *p<0.05, **p<0.01,***p<0.001,****p<0.0001 Tukey’s multiple comparisons test, n≥9 guts/sample.

In order to assess the effects of Sob at the cellular level we drove sob knockdown and overexpression using a temperature inducible ISC/EB specific lineage tracing system^30^. Knockdown of *sob* in young flies resulted in an increase in the proportions of labelled cells indicative of enhanced stem cell proliferation (Figure 3A-C). Consistent with these findings sob knockdown with the ISC/EB geneswitch driver increases the mitotic index of the midgut in young flies (Supplementary Figure S3A) .In addition, staining for the enteroendocrine (EE) cell marker prospero found a reduced proportion of EE cells (Figure 3D). Overexpression of sob did not impact the proportion of labelled cells, consistent with the finding that Sob overexpression had no effect on the mitotic index in young flies but did increase the proportion of EE cells in the midgut (Figure 3). Similar results were obtained using a temperature inducible ISC/EB driver to label just the ISCs and EBs without lineage tracing (Supplementary Figure S3C-G). Collectively these results suggest Sob constrains ISC proliferation and contributes to maintaining the balance of EC: EE differentiation during normal homeostasis.

**Figure 3:**
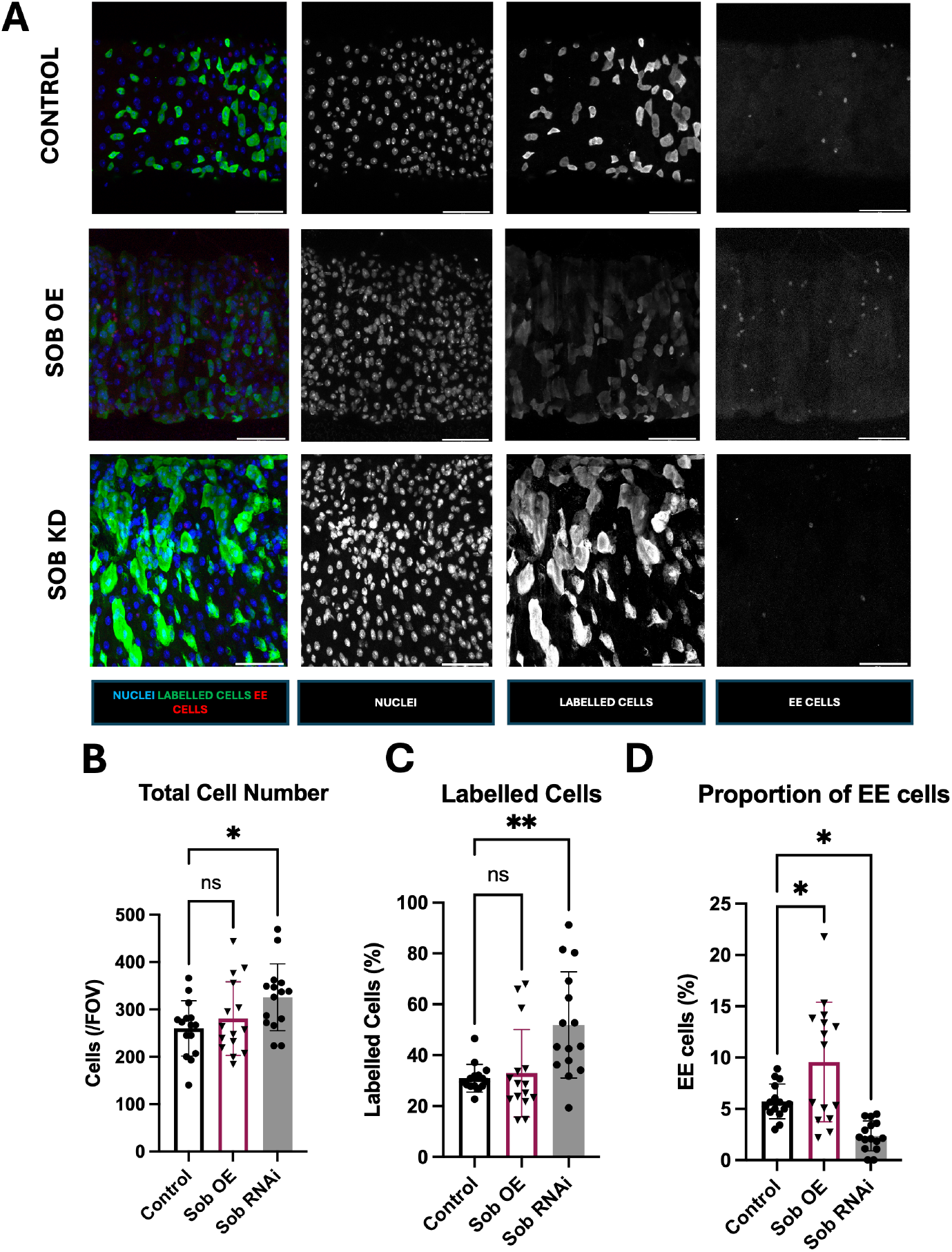
Stem/ progenitor Sob Regulates ISC Proliferation and Fate. (A) Representative maximum intensity-projected confocal z stacks (brightness and contrast enhanced for clarity) of control, sob knockdown and sob overexpression. Greyscale images show nuclei (DAPI), labelled cells (esgTSFlip), and EE cells (pros). Merged images show nuclei (blue), ISCs/EBs (green), and EE cells (red). Scale bars = 50 µm. Quantification of (B) cell number per field of view (FOV), (C) proportion of labelled cells, (D) proportion of EE cells.*p<0.05, **p,0.01, ****p<0.0001 Ordinary one- way Anova with Dunnett’s multiple comparisons, n≥15 guts.

### Sob regulates transcriptional regulators and signalling pathway components

We next aimed to identify the regulators acting downstream of Sob. RNAseq was performed on whole midguts in which sob was knocked down or overexpressed specifically in the ISC/EBs. Using stringency cut offs of adjusted p-value <0.05, log2 fold change > or < 0 to identify initial candidate lists, knockdown of sob led to downregulation of 357 genes and upregulation of 73 genes, while overexpression led to downregulation of 888 genes and upregulation of 1166 genes (Figure 4A and B; Supplementary Figure S4; Supplementary Tables). 80 genes showed reciprocal regulation including the transcription factor Exex which has been described as a marker of EE cells in a specific region of the midgut^31^. *exex* is upregulated on *sob* knockdown, when EE numbers are decreased, and downregulated on Sob overexpression, when EE numbers are increased. An exex-GFP protein trap line validated the regulation of *exex* by Sob showing upregulation on *sob* knockdown and downregulation on Sob overexpression (Figure 4E and F).

**Figure 4:**
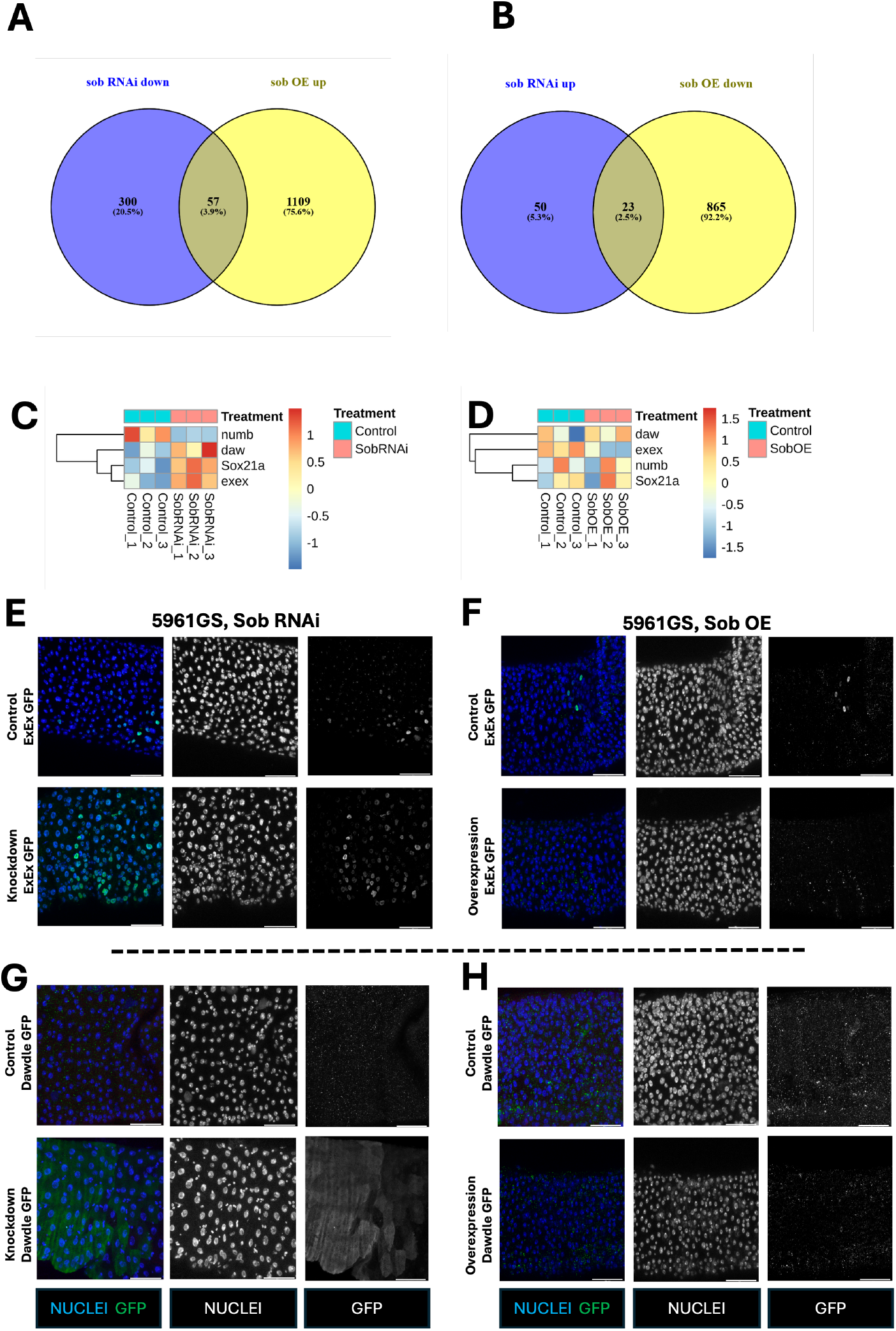
Transcriptomic changes on sob knockdown and overexpression. (A,B) Venn diagrams of RNAseq data showing differential gene expression in the adult midgut following sob RNAi or sob overexpression in stem and progenitor cells. n=3 biological replicates. (C,D) Heatmaps showing expression of candidate targets in sob knockdown (C) and overexpression (D). (E-H) Representative maximum intensity-projected confocal z stack images (brightness and contrast enhanced for clarity) of exexGFP (E and F) and dawGFP (G and H) in sob knockdown (E and G) and overexpression (F and H) and respective controls. Greyscale images show nuclei (DAPI), and reporter (GFP). Merged images show nuclei (blue) and reporter (green) . Scale bars = 50 µm.

Focussing on signalling pathways that maintain intestinal homeostasis and differentiation and may act downstream of Sob we identified Dawdle and Numb as candidates. The TGF-ligand Dawdle, which has been shown to promote EB to EC differentiation^32^, is upregulated on sob knockdown. The negative regulator of Notch signalling Numb has an important role in EE differentiation^33^ and was downregulated on *sob* knockdown. Upregulation of a dawdle GFP reporter on sob knockdown (Figure 4G, H) and antibody staining for Numb protein (Supplementary Figure S4) validated these signalling factors as potential Sob targets. Given that knockdown of *sob* resulted in reduced *numb* expression we hypothesised that Sob may negatively regulate Notch signalling via Numb. A Notch pathway transcriptional reporter (Su(H)GBE-nlsGFP) showed increased expression on *sob* knockdown and reduced expression on Sob overexpression (Figure 5).

**Figure 5:**
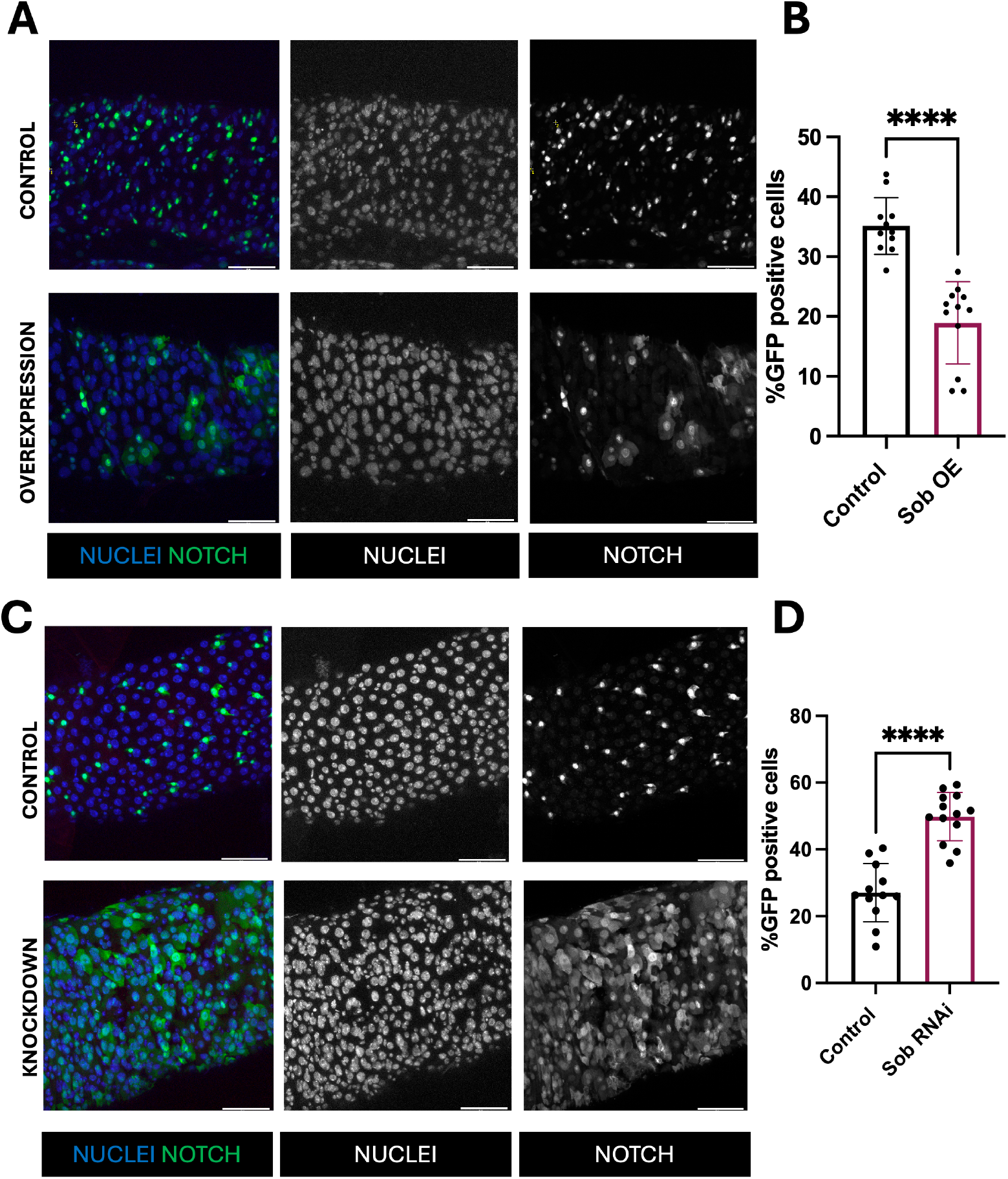
Sob enhances Notch signaling. (A,C) Representative maximum intensity-projected confocal z stacks (brightness and contrast enhanced for clarity) of sob overexpression (A) and knockdown (C) and respective controls. Greyscale images show nuclei (DAPI), and Notch transcriptional reporter (Su(H)GBE-GFPnls). Merged images show nuclei (blue) and Notch reporter activity (green). Scale bars = 50 µm. (B,D) Quantification of Notch reporter positive cells (notch positive) in sob overexpression (B) and knockdown (D). *p<0.05, **p,0.01, ****p<0.0001 unpaired t test, n≥15.

### Dawdle, Numb and Exex Function Downstream of Sob

Our results suggest that Exex, Dawdle and Numb could function downstream of sob to regulate proliferation and differentiation. To test this hypothesis, we conducted a series of experiments to assess whether they could rescue or supress *sob* phenotypes. Our expression analysis suggests that *exex* is negatively regulated by Sob and upregulated when *sob* is knocked down. We therefore knocked down *exex* in a *sob* knockdown background and saw suppression of *sob* knockdown induced proliferation and EE cell loss (Figure 6A-E). Reciprocally overexpression of Exex in a Sob overexpression background suppressed the increase in EE cell numbers and caused overproliferation (Figure 6F-J). *dawdle*, which shows a similar pattern of regulation to exex was also knocked down in a sob knockdown background. *Dawdle* knockdown has no impact on the proliferation phenotype but did rescue EE cell numbers (Figure 6A-E). Expression profiling suggested Sob may positively regulate Numb as numb expression is reduced in sob knockdown, we therefore knocked down Numb in a Sob overexpression background and saw a suppression of the increase in EE cells but no effect on proliferation (Figure 6F-J). Collectively these results show that Sob’s effect on proliferation and differentiation are mediated by parallel downstream mechanism involving exex, Numb and dawdle. Exex is involved in both proliferation and differentiation effects whereas Dawdle and Numb only affect differentiation (Figure 7).

**Figure 6:**
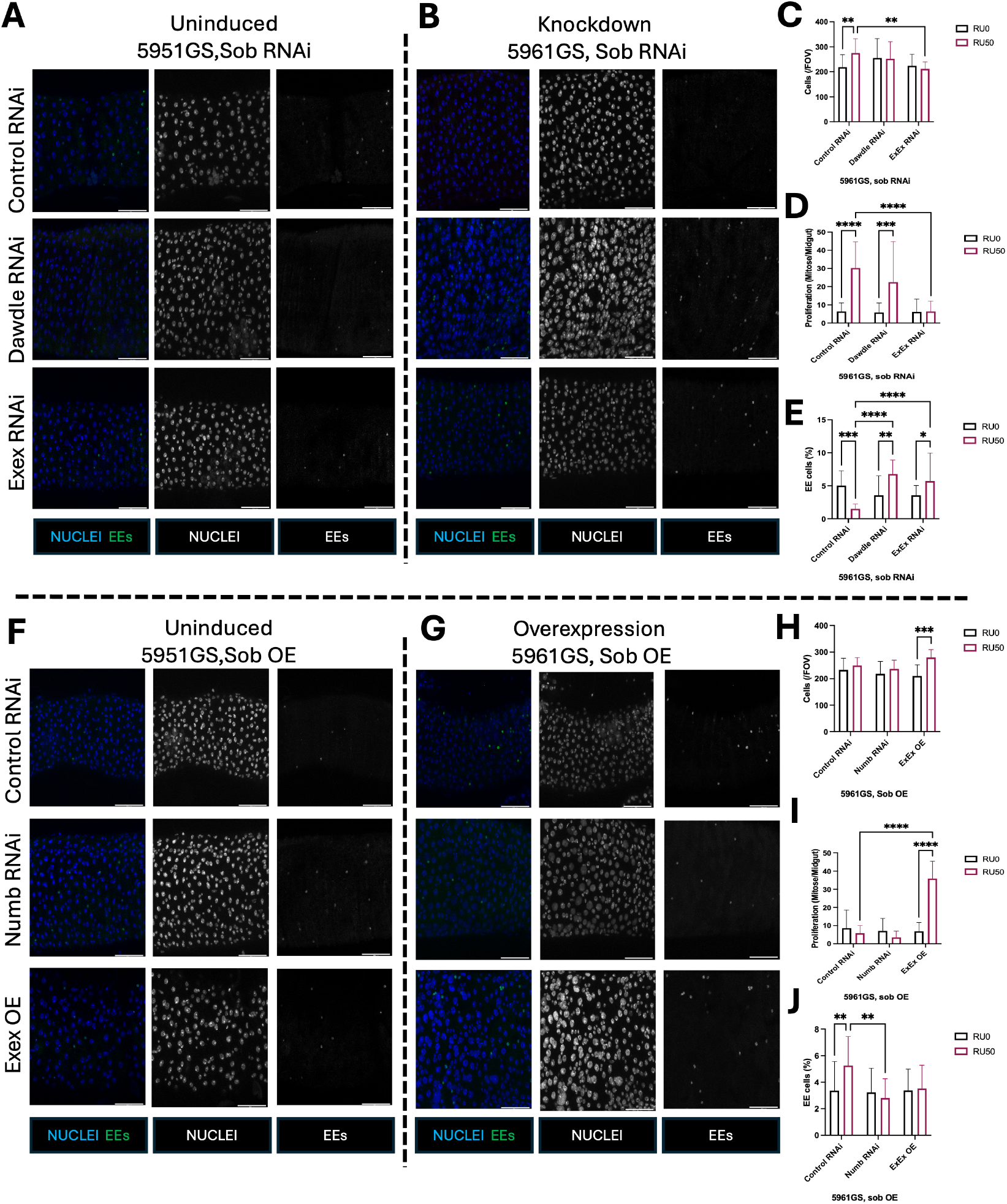
daw, numb and exex function downstream of Sob. (A,B,F,G) Representative maximum intensity-projected confocal z stacks (brightness and contrast enhanced for clarity) of sob co-knockdown with dawdle or exex and respective controls (A and B). and sob overexpression with Numb RNAi or Exex OE and respective controls (F and G). Greyscale images show nuclei (DAPI) and EEs (pros) Merged images show nuclei (blue) and EEs (green). Scale bars = 50 µm. Quantification of total cell number (C, H) Proliferation (D,I) and % EEs (E,J). *p<0.05, **p,0.01, ****p<0.0001 Mixed effects model with Sidak’s multiple comparisons, n≥18 midguts.

**Figure 7:**
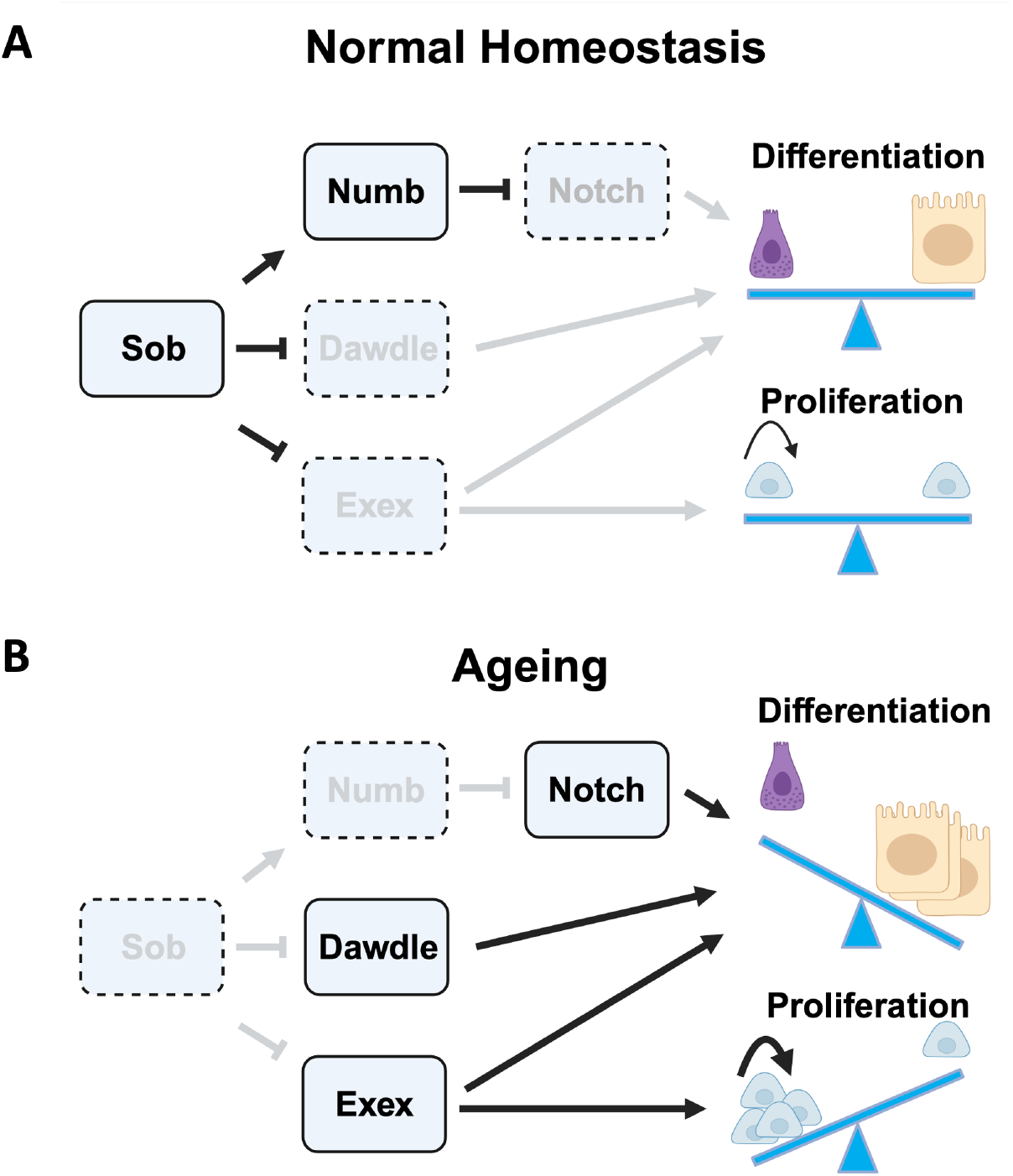
Model of Sob Function in Homeostasis and ageing. Representation of Sob function in homeostasis (A) and ageing (B). Arrows indicate activation, lines with bars indicate repression. Bold protein names in solid boxes indicates expression, Grey names in dashed boxes indicate loss or reduction in expression. Created in BioRender. DOUPE, D. (2026) https://BioRender.com/gdijlg7.

## Discussion

The Odd skipped family of transcription factors including Sob are conserved from flies to humans and have well characterised roles in a range of developmental contexts^34-38^. However, their function in adult tissue homeostasis and ageing have not been well explored. Here we have shown that family member Sob is expressed in *Drosophila* ISCs and EBs and regulates balanced proliferation and differentiation to maintain homeostasis. Its expression is lost with age and overexpression extends median lifespan. The mammalian Sob ortholog OSR2 shows similarities in expression in mouse ISCs and TA cells and is similarly lost with age correlating with a possible conserved function.

### Sob as a key component of the transcriptional regulation of homeostasis

Previous work on Sob showed roles in development including in leg joints, where it interacts with Notch^34^, and in the early eye disc^35^. Its identification as a regulator of intestinal stem cell fate adds it to a growing network of transcription factors, many of which show some conservation of function to mammalian systems. There include Escargot, SNAI1, Sox21a, Sox100B and the E(spl) complex genes and chromatin regulators such as Chronophage^9-22^. Overexpression of some homeostatic transcriptional regulators such as Caudal and Chronophage have been shown to enhance age-related dysplasia^22,24^. This opposing role to Sob is consistent with their homeostatic function in promoting stem cell maintenance and proliferation. Changes in transcription factor expression also have significant effects in the mammalian intestine. TFs regulating genes involved in ISC proliferation and differentiation that show changes in expression with age affect ISC function as assayed by ability to generate organoids *in vitro*^39^.

Given that Sob impacts both stem cell homeostasis and ageing at the organismal level it is not surprising that it targets important regulators of major niche signaling pathways. Mis-regulated signalling in the stem cell niche plays a major role in ageing. Overactivation of pathways such as JAK/STAT and EGFR/Ras drives excessive ISC proliferation^40,41^. Reduced Notch signalling impairs EB differentiation into ECs leading to accumulation of undifferentiated progeny^42^. Consistent with their reciprocal regulation by Sob, Numb and the TGF-ß ligand Dawdle have distinct roles in regulating the balanced differentiation of ISC progeny. Numb has previously been shown to be asymmetrically segregated on ISC division and play an important role in promoting EE fate specification^17,33,43^. Dawdle has been shown to promote EB to EC differentiation through its receptor Babo-C with a role in regulation of gut size in response to nutrient availability^32^. Dawdle and Numb’s functions downstream of Sob are consistent with misregulation of these known roles. A recent study demonstrated a link between Numb and BMP signalling in the midgut epithelium with epithelial BMP signalling working in parallel with Numb to constrain Notch signalling and maintain homeostasis^44^. Sob’s promotion of Numb expression may reinforce this mechanism and given the often-complex relationship between Activin and BMP branches of TGF-superfamily signalling^45^ it is possible that the repression of daw also contributes.

We also show that the transcription factor Exex is an important target of Sob. Exex has previously been identified from single cell RNA sequencing experiments as a region-specific marker of EE cells^31^ that regulates expression of specific peptide hormones. Our findings suggest Exex may itself have an unstudied functional regulatory role in homeostasis that warrants further research. In addition, while we have shown Notch signalling, Dawdle and Exex play key downstream roles it is likely other mechanisms function downstream of Sob including possible non-autonomous effects, which could be induced by Dawdle or other secreted targets.

### Sob / OSR2 as a conserved switch in ISC ageing

Similarly to Sob, OSR2 has been shown to have a range of roles during development including in limb and craniofacial development and in osteogenesis^36-38,46^. Its expression has been reported in the developing mouse intestine^47^ and association with certain tumours such as invasive bladder cancer and endometrial cancer may point to a role in regulation of proliferation or differentiation^48,49^. However, its function in normal adult tissues is less well explored. The similarities in OSR2’s expression pattern to Sob in homeostasis and reduced expression with age raise the possibility of a conserved role for Sob / OSR2 in mammalian intestinal stem and TA cells. In addition, the expression of OSR2 in Goblet cells may suggest a related role in differentiation to that of Sob. Notch signalling plays a critical role in Goblet cell differentiation^50,51^ and Exex ortholog Mnx is enriched in goblet cells (proteinatlas.org)^27,28^. This correlates with a potential conserved role of MNX and Notch acting downstream of OSR2 as Exex and Numb act downstream of Sob. Further studies will be needed to explore the functional role of OSR2 in mammalian intestine such as loss of function studies *in vivo* or in organoid cultures, and any conserved relationship with MNX and Notch signalling.

Overall, our findings position Sob as key switch in intestinal stem cell regulation, not only ensuring maintenance of epithelial homeostasis but also healthy organismal ageing. The ability of ISC/EB Sob expression levels to impact ageing at the organismal level adds to a growing body of evidence that intestinal and stem cell homeostasis is critical to ageing and health and that strategies targeting stem cells can delay ageing^40,52^.

## Materials and Methods

### Drosophila Stocks and Genetics

Full details of stocks used are listed in Supplementary Table 1. Flies were maintained on standard cornmeal medium (1% agar, 3% inactivated yeast, 1.9% sucrose, 3.8% dextrose, and 9.1% cornmeal; see Supplementary Table 2 for reagents) at 18 °C, 60% humidity with 12-h light/dark cycle and transferred to fresh food every three weeks.

Experimental crosses were maintained at 25 °C unless otherwise stated. Progeny were collected, aged to the required time points, and used as mated females for all experiments. For temperature-inducible experiments, crosses were maintained at 18 °C, progeny were and aged for 7 days at 18 °C and then shifted to 29 °C for a further 7 days to induce transgene expression. RU486-inducible expression was achieved by feeding cornmeal medium supplemented with 50 µg/mL mifepristone (RU486; Cayman Chemical, 10006317) or vehicle control (ethanol) for the indicated durations.

### Midgut Immunofluorescence and Confocal Microscopy

Whole adult female guts were dissected in cold phosphate-buffered saline (PBS) and fixed in 4% paraformaldehyde for 30 min at room temperature. Samples were blocked in PBS containing 1% bovine serum albumin, 0.5% Triton X-100, and 5% normal goat or donkey serum according to the secondary antibody. Primary antibodies (Supplementary Table 3) were incubated overnight at 4 °C with rocking. After washing, samples were incubated with fluorophore-conjugated secondary antibodies (Supplementaty Table 3) were incubated for 2 h at room temperature. Midguts were then counterstained with DAPI, and mounted in Vectashield.

Images of the posterior midgut were acquired using a Leica LSM SP5 confocal microscope with a 63x objective (0.88 µm z-step, 1024 × 1024 pixels). Acquisition parameters were kept constant within each experiment. Image analysis was performed using Fiji (ImageJ). Maximum-intensity projections were used for quantification, and representative images were brightness-adjusted where indicated. Statistical analyses were performed using GraphPad Prism 10. Outliers were removed using the ROUT method, and data normality was assessed using Shapiro–Wilk or Kolmogorov–Smirnov tests. Significance was determined using unpaired t-tests or Mann–Whitney U tests for two groups, and one-way ANOVA or Kruskal-Wallis tests for multiple groups. Significance thresholds were defined as p ≤ 0.05 (*), p ≤ 0.01 (**), p ≤ 0.001 (***), and p ≤ 0.0001 (****).

### Mouse Samples and Immunohistochemistry

Paraffin-embedded intestinal tissues from age-matched young and old mice were obtained from Newcastle University. All procedures complied with the Animals (Scientific Procedures) Act 1986 and EU Directive 2010/63/EU, and were approved by the UK Home Office (PPL P3052AD70) and Newcastle University Animal Welfare Ethical Review Board (AWERB 425). Mice were housed in same-sex groups under a 12-h light/dark cycle at 20 ± 2 °C and monitored daily until study endpoints.

Immunohistochemistry (IHC) was performed on paraffin-embedded intestinal sections to detect OSR2 expression. Sections were incubated at 60 °C for 45 min, deparaffinised in Histoclear, rehydrated through graded ethanol, and immersed in distilled water. Antigen retrieval was carried out in 10 mM citrate buffer (pH 6.0) using a pressure cooker. Sections were washed in Tris-buffered saline with 0.1% Tween-20 (TBST) and treated with 0.3% hydrogen peroxide in methanol for 10 min to quench endogenous peroxidase. After blocking in 10% normal goat serum (NGS) in TBST for 1 h at room temperature, sections were incubated overnight at 4 °C with rabbit anti-OSR2 antibody (Supplementary Table 3) at 1:250 dilution. The next day, sections were washed in TBST and incubated with the Vector ImmPRESS HRP anti-rabbit polymer reagent for 30 min. Staining was developed using SigmaFAST DAB for 5-10 min, followed by haematoxylin counterstaining, bluing in Scott’s tap water, dehydration through graded ethanol, clearing in Histoclear, and mounting in DPX (BDH Laboratory Supplies). Slides were scanned using an Aperio Slide Scanner (Leica Biosystems), and images were processed and analysed in QuPath.

### RNA Sequencing

Total RNA was extracted from dissected midguts (10 per sample, 3 biological replicates per condition) using the Monarch Total RNA Miniprep Kit (NEB, T2010S; see Supplementary Table 2). RNA quality was confirmed by NanoDrop (A260/A280 > 2.0, A260/A230 > 2.0). Poly(A)-enriched libraries were prepared and sequenced by Novogene on the Illumina NovaSeq 6000 platform (PE150, ∼20 million reads/sample).

Raw reads were processed with fastp for adapter trimming and aligned to the *Drosophila* melanogaster BDGP6.46 genome to obtain gene level counts using STAR (v 2.7.11b)^53^ and corresponding gtf file (https://ftp.ensembl.org/pub/release-112/gtf/drosophila_melanogaster/Drosophila_melanogaster.BDGPG.4G.112.gtf.gz). Differential expression was analysed using DESeq2 (v 1.44.0)^54^. Heatmapswere generated in R using pheatmap (v 1.0.12)^55^ Datasets will be made available from GEO (GSE330270).

### RT-qPCR

RNA was extracted from 10 dissected midguts per sample as described above. RNA (1 µg) was reverse-transcribed using the Ultrascript 2.0 cDNA Synthesis Kit (PCR Biosystems). Quantitative PCR was performed with Power SYBR Green PCR Master Mix (Applied Biosystems) on a Bio-Rad CFX Connect system. Reactions (10 µL) contained 5 µL SYBR mix, 0.5 µL of each primer (10 µM), 1 µL cDNA, and 3.5 µL RNase-free water. Primer sequences (Table 2.5) were obtained from FlyPrimerBank^56^.

Cycling conditions were: 95 °C for 10 min, followed by 40 cycles of 95 °C for 15 s, 60 °C for 1 min, and 70 °C for 30 s. Melt-curve analysis confirmed specificity. Relative expression was calculated using the ΔΔCt method and normalised to Actin or GAPDH. Each condition included five to six biological replicates analysed in technical triplicates. Statistical analyses were performed as described above.

## Acknowledgements

FS was supported by NLD (Newcastle-Liverpool-Durham) BBSRC Doctoral Training Partnership 3 (BB/T008695/1). LCG is supported by Cancer Research UK (DRCPFA-Nov22/100001), The Medical Research Council (MC_PC_21046 and MC_PC23036), The LifeArc centre for Rare Mitochondrial Diseases (REF:10748), and the NIHR Newcastle Biomedical Research Centre. Stocks obtained from the Bloomington Drosophila Stock Center (NIH P40OD018537) were used in this study. We are grateful to Rebecca Clark, Norbert Perrimon, Lucy O’Brien and Jens Januschke for providing fly stocks and antibodies used in this study; and to Rebecca Clark, Patricia Muller and Alistair McGregor for helpful comments on the manuscript.

## Author Contributions

FS – conceptualization, investigation, data analysis, data presentation, writing – original draft, review and editing. WW – data curation, data analysis, data visualisation. AS – supervision, investigation, data analysis . LG – supervision, funding acquisition, writing – review and editing. DD – conceptualization, funding acquisition, investigation, supervision, writing – original draft, review and editing

