## Supplementary Figures and Tables for "The Transcription Factor Sob Maintains Intestinal Stem Cell Homeostasis to Delay Ageing"

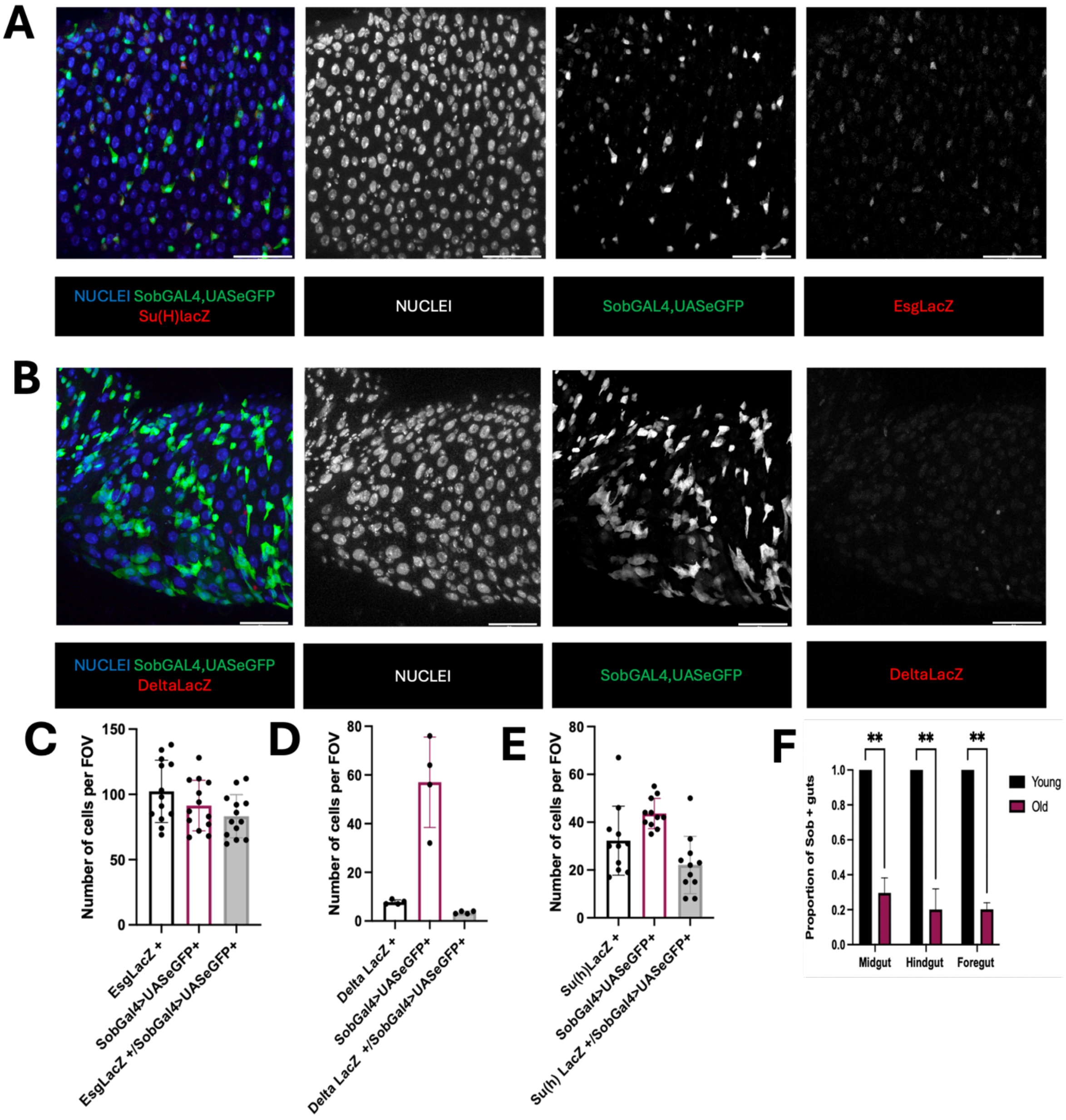


**Supplementary Figure S1: OSR2/Sob expression in ISCs.**

Representative maximum intensity-projected confocal z stack images with brightness and contrast uniformly enhanced for clarity. The images show the posterior midgut of 10-day old sob-Gal4,UASeGFP with Su(h)lacZ (A) and Delta (B). Grey-scale images show nuclei (DAPI), sob expressing cells (sob-GAL4, UASeGFP), and ISC/EBs (LacZ). Merged images show nuclei (blue), sob (green), and ISC/EBs (red). Scale Bar = 50µm. Bar Graphs showing the number of sob-expressing cells (sob-GAL4>EGFP) overlapping with esg+ ISC and EBs (C), Su(H)GBE-lacZ+ (EBs) (D), or Delta+ (ISCs) (E). Proportion of guts with positive sob expression: bar chart to show how the expression pattern of sob changed with age in young homeostatic midguts versus old midguts (F). $^{**}p<0.01$

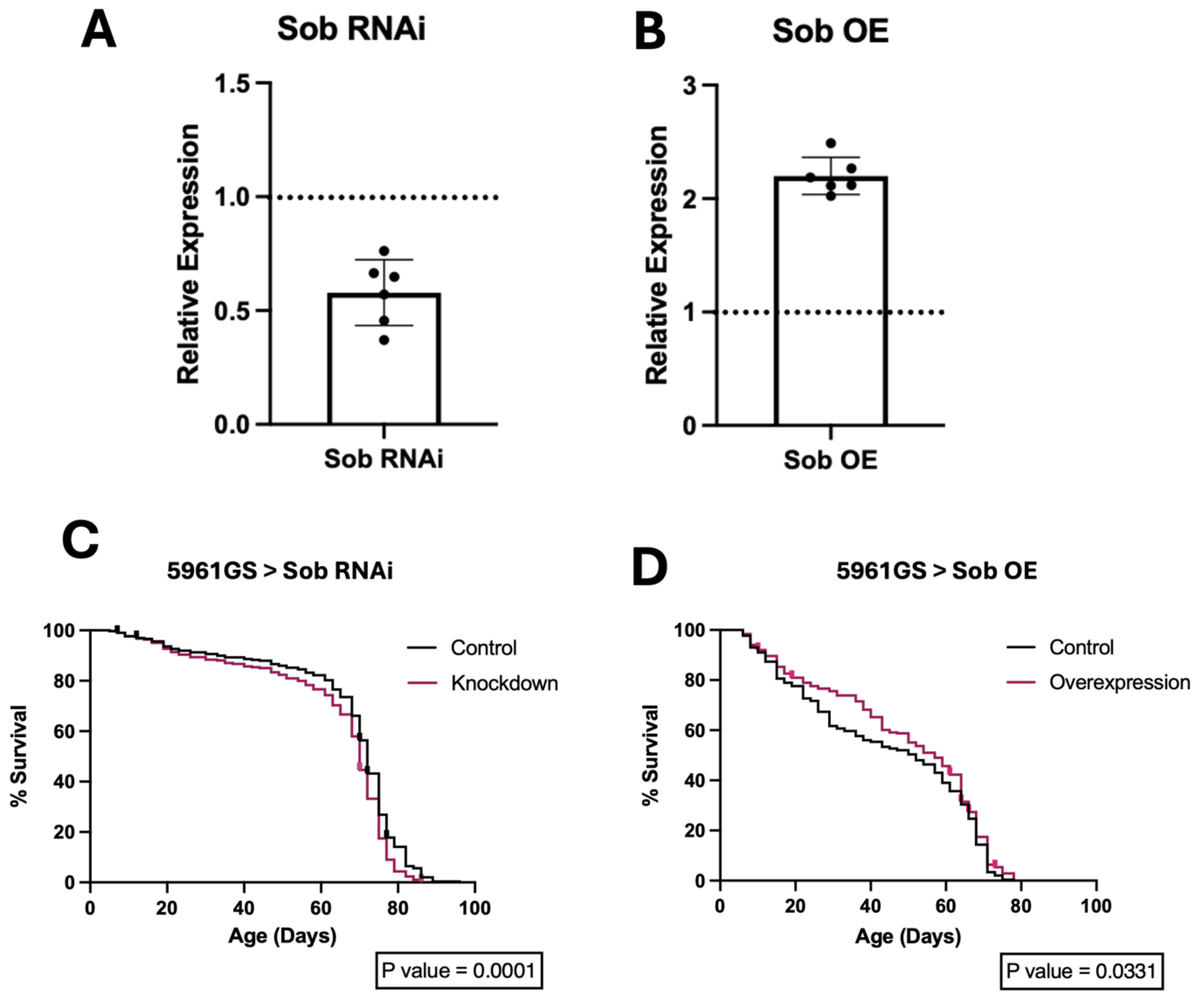


**Supplementary Figure S2: Sob Impacts Lifespan.**

Changes in sob expression levels in whole gut samples using the ubiquitous driver DaGS with specified RNAi (A) and overexpression (B) lines, relative to GAPDH. The bars show the fold change. Calculated using an unpaired t test. n=10 guts/sample, ≥5 replicates. **p < 0.01, *** p < 0.001, ****p < 0.0001. Lifespan graphs showing second replicate of sob knockdown with sob RNAi (C) and overexpression (D) using the driver 5961GS. Lifespans were tested with log-rank (Mantel-Cox) test, p values are indicated on each plot. n=300 flies/condition.

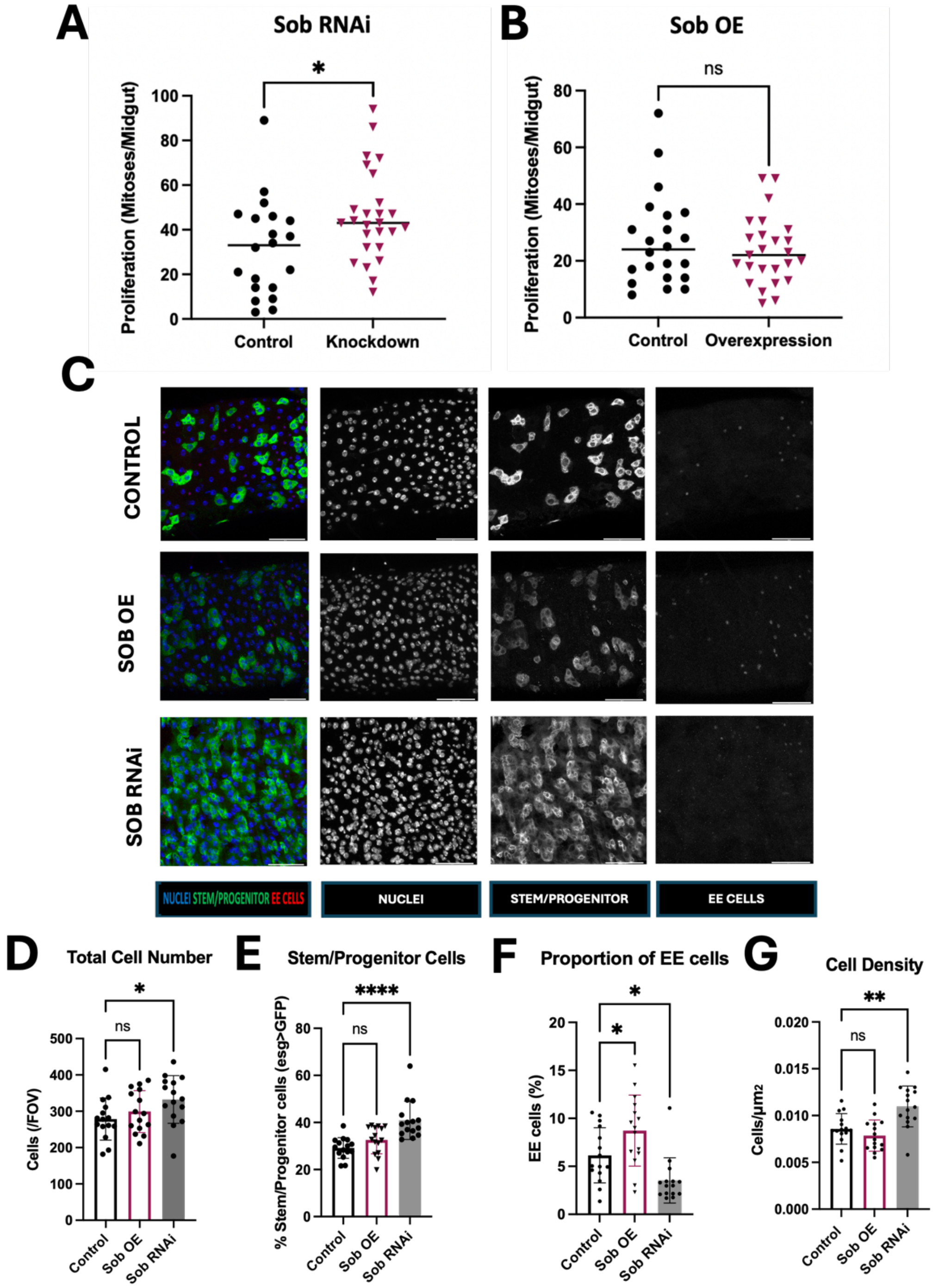


### Supplementary Figure S3: Sob Regulates ISC EBs.

A significant increase in mitotic cells was observed following knockdown of sob in young flies (A). No significant change was observed on overexpression in young flies (B). Outliers were identified using the ROUT method, followed by an unpaired t test n≥10 guts from 3 separate replicates. No* = ns, *p<0.05, **p,0.01, ****p<0.0001. (C)Representative maximum intensity-projected confocal z stack images with brightness and contrast enhanced for clarity of control RNAi, sob knockdown and sob overexpression. Grey- scale images show cells per field of view Nuclei (DAPI), ISCs/EBs (esg$^{ts}$GFP), and EE cells (pros+). Merged images show nuclei (blue),

ISCs/EBs (green), and EE cells (red). representative of n≥20 across 3 biological replicates. Scale bars = 50 µm. Changes in cell number per field of view (D), the proportion of stem/progenitor cells (E), the proportion of EE cells (F) and the cell density (G) have been quantified in knockdown and overexpression backgrounds. Calculated using an Ordinary one-way Anova with multiple comparisons. n≥15 guts across 3 biological replicates. No* = ns, *p<0.05, **p,0.01, ****p<0.0001.

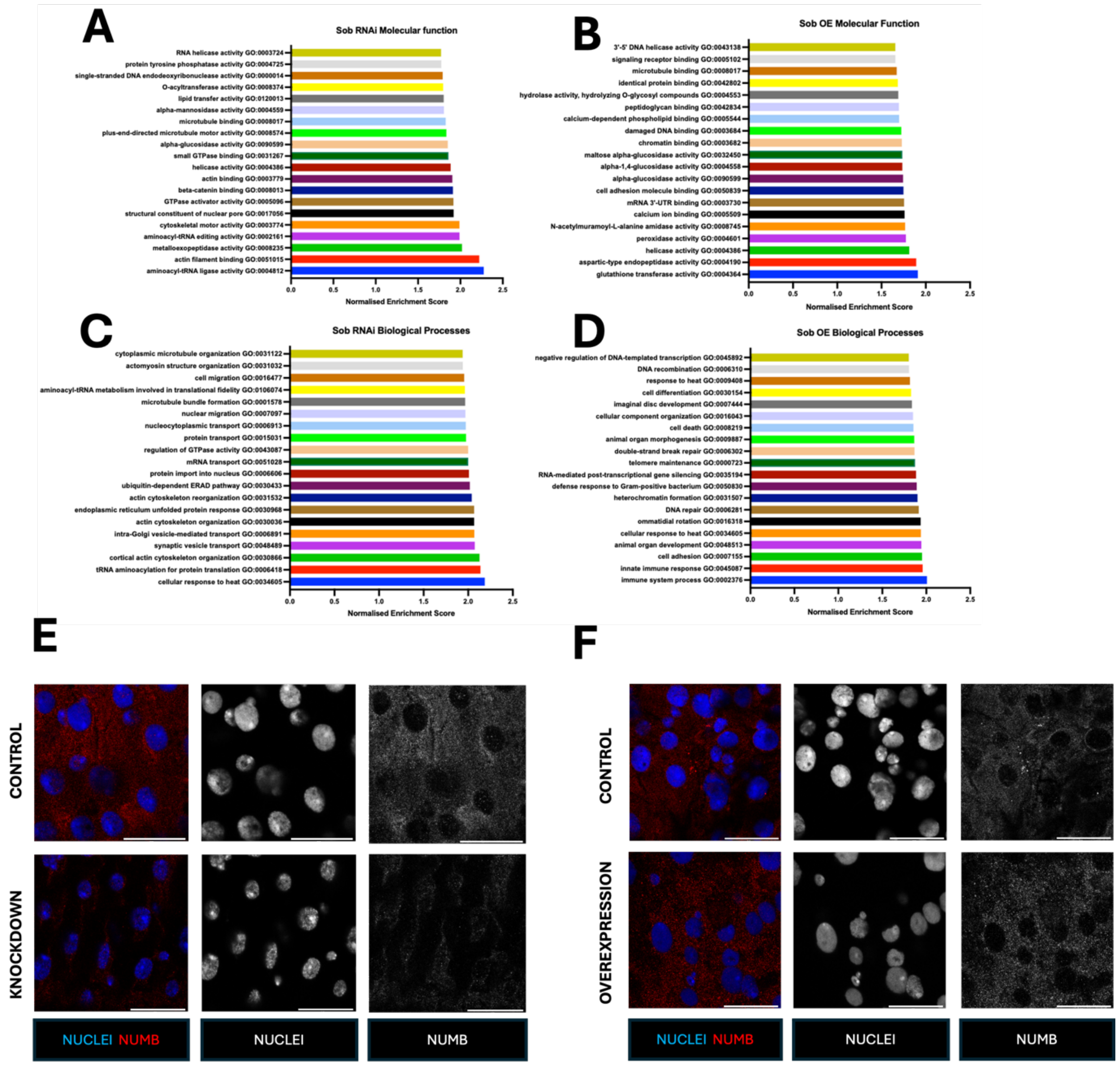


**Supplementary Figure S4: Sob regulated genes.**

GO showing molecular function (A,B) and biological processes (C, D) in sob knockdown and overexpression backgrounds. Top 20 most enriched categories are shown. Representative maximum intensity-projected confocal z stack images with brightness and contrast enhanced for clarity of numb (red) in sob overexpression/knockdown (E,F) and respective controls. Grey- scale images show cells per field of view Nuclei (DAPI),  and Numb (red). Merged images show nuclei (blue) and numb (red). Representative of n≥10 across 3 replicates. Scale bars = 25 µm.

### Supplementary Table 1: Fly Stocks

| Line | Source Ref | Genotype |
|---|---|---|
| UAS-eGFP | BDSC_6874 | *w[*]; P{w[+mC]=UAS-2xEGFP}AH2* |
| 5961 GeneSwitch | 5961 (O'Brien lab) | *5961-GeneSwitch-Gal4* |
| esg$^{TS}$Flip | Perrimon Lab | *esg$^{TS}$-GFP-Flp-Out* |
| Luciferase | 31603 | *y[1] v[1]; P{y[+t7.7] v[+t1.8]=TRiP.JF01355}attP2* |
| esg$^{TS}$GFP | Perrimon lab | esgGAL4,UAS-GFP,tubGAL80ts |
| esg-lacZ | Perrimon lab | y[1] w[67c23];P{w[+mC]=lacW}esg[k00606]/CyO |
| w1118;If/SM6a;TM2/TM6c,Sb | Clark lab | w1118;If/SM6a; TM2/TM6c,Sb' |
| esgGal4 | BDSC_93857 | *y[1] w[*]; P{w[+mW.hs]=GawB}NP5130* |
| Sob GFP | BDSC_92326 | *y[1] w[*]; PBac{y[+mDint2] w[+mC]=sob GFP.FPTB}VK00037* |
| Sob Gal4 | BDSC_83247 | *y[1] w[*]; TI{GFP[3xP3.cLa]=CRIMIC.TG4.2}sob[CR01007-TG4.2]/SM6a* |
| Sob RNAi | BDSC_34648 | *y[1] sc[*] v[1] sev[21]; P{y[+t7.7] v[+t1.8]=TRiP.HMS01123}attP2* |
| UAS Sob | BDSC_7085 | *w[*]; P{w[+mC]=UAS-sob.G}15.1* |
| Dawdle Mimic | BDSC_43001 | *y[1] w[*]; Mi{y[+mDint2]=MIC}daw[MI05383]* |
| Exex GFP | BDSC_66749 | *w[1118]; P{y[+t7.7] w[+mC]=exex-GFP.FPTB}attP40* |
| Bmm GFP | BDSC_94600 | *w[*]; TI{TI}bmm[GFP]* |
| Sima GFP | BDSC_43957 | *w[1118]; PBac{y[+mDint2] w[+mC]=sima-GFP.B.FPTB}VK00037* |
| Bgm LacZ | BDSC_28120 | *w[*]; P{w[+mC]=lacW}bgm[1]* |
| SobGal4,UASeGFP/cyo | This study | *SobGal4,UASeGFP/cyo* |
| 5961GS/cyo: Sob RNAi/TM6 | This study | *5961GS/cyo: Sob RNAi/TM6* |
| 5961GS/cyo:Sob OE/TM6 | This study | *5961GS/cyo: Sob OE/TM6* |
| DaGS | Rebecca Clark | |
| If/cyo: SobRNAi | This study | *If/cyo: SobRNAi* |
| If/cyo: SobOE | This study | *If/cyo: SobOE* |
| SobGal4/ cyo: MKRS/TM6 | This study | *SobGal4/ cyo: MKRS/TM6* |
| If/cyo | Clark Lab | *If/cyo* |
| 5961GS: TM2/ TM6 | Doupe Lab | *5961GS: TM2/ TM6* |
| TM2/TM6 | | *TM2/TM6* |
| Dawdle RNAi | BDSC_34974 | *y[1] sc[*] v[1] sev[21]; P{y[+t7.7] v[+t1.8]=TRiP.HMS01110}attP2* |
| Sima OE | BDSC_9582 | *w[*]; P{w[+mC]=UAS-sima.B}2* |
| Sima RNAi | BDSC_26207 | *y[1] v[1]; P{y[+t7.7] v[+t1.8]=TRiP.JF02105}attP2* |
| Bmm RNAi | BDSC_25926 | *y[1] v[1]; P{y[+t7.7] v[+t1.8]=TRiP.JF01946}attP2/TM3, Sb[1]* |
| Bmm OE | BDSC_76600 | *w[*]; P{w[+mC]=UAS-bmm.cGa}2* |

| | | |
|---|---|---|
| Bgm RNAi | BDSC_28639 | *y[1] v[1]; P{y[+t7.7] v[+t1.8]=TRiP.JF03054}attP2/TM3, Sb[1]* |
| Exex RNAi | BDSC_57709 | *y[1] sc[*] v[1] sev[21]; P{y[+t7.7] v[+t1.8]=TRiP.HMC04898}attP2* |
| Exex OE | BDSC_9929 | *w[*]; P{w[+mC]=UAS-exex.B}H1/CyO* |
| Numb RNAi | BDSC_35045 | *y[1] sc[*] v[1] sev[21]; P{y[+t7.7] v[+t1.8]=TRiP.HMS01459}attP2* |

### Supplementary Table 2: Reagents

| Product | Supplier | Catalogue Number |
|---|---|---|
| Phosphate Buffered Saline Tablets 0.01M | Fisher Bioreagents | BP2944100 |
| Bovine Serum Albumin | Fisher Scientific | BP-1600-100 |
| Vectashield Antifade Mounting Medim | 2B Scientific | H-1000 |
| Triton X-100 | Sigma Aldrich | 9002-93-1 |
| Apex *Drosophila* Agar Type II (pH = 6.13) | SLS | #AN-171047 |
| Inactive Dry Yeast) | SLS | FLY#62-106 |
| Sucrose Crystallized | Duchefa Biochemie | S0809.5000 |
| D-(+)-Glucose Anhydrous (MW = 180.16) | Melford | G32040, Lot. 42613-42844 |
| Cornmeal Yellow | SLS | FLY#62-100 |
| Mifepristone – Cayman Chemicals | Cambridge Bioscience | CAY10006317 |
| Phosphoric acid | VWR | 20624.262 |
| Propionic acid | Sigma | P5561 |
| Nipagin | SLS | FLY1136 |
| Optically Clear Adhesive Seal Sheets | ThermoFisher Scientific | AB-1170 |
| Water (for RNA work), DEPC-treated and nuclease free, autoclave | Fisher Bioreagents | BP561-1, Lot. 207040 |
| Ethanol, molecular grade | Fisher Bioreagents | BP2818-500 |
| EDTA | Thermofisher | EN0521 |
| Paraformaldehyde (PFA), 16%, 2x10mL | Fisher | 11490570 |
| Normal Goat Serum (NGS), 2mL | Gibco | PCN5000 |
| InvitrogenTM DAPI (4',6- Diamidino-2-Phenylindole, Dihydrochloride) | Fisher Scientific | 10184322 |
| Parafilm M Sealing Film | SLS | FIL1024 |
| DEPC- Rnase free Water | Fisher Bioreagents | BP561-1 |
| Normal Donkey serum (NDS) | Sigma Aldrich | D9663 |
| ImmPRESS® HRP Goat Anti-Rabbit IgG Polymer Detection Kit, Peroxidase | Vector Labs | MP-7451 |
| 3,3′-Diaminobenzidine (DAB) | Sigma Aldrich | D4293-5SET |
| Monarch Total RNA Miniprep Kit | NEB | T2010S |
| Ultrascript 2.0 cDNA Synthesis Kit | PCR Biosystems | PB30.31-10 |
| Power SYBR Green PCR Master Mix | Applied Biosystems | 4367659 |

### Supplementary Table 3: Antibodies

| Type | Species | Antibody | Concentration | Supplier | Reference |
|---|---|---|---|---|---|
| Primary | *Drosophila* | Anti-Prospero Mouse IgG Monoclonal Antibody | 1:100 | Developmental Studies Hybridoma Bank (DSHB) | MR1A |
| Primary | *Drosophila* | Anti-phospho-histone-H3 (Ser10), Mitosis Marker Rabbit Polyclonal Antibody | 1:1000 | Merck Life Science | 06-570 |
| Primary | *Drosophila* | Anti-Green Fluorescent Protein (GFP) Chicken IgY Polyclonal Antibody | 1:2000 | Abcam | ab13970 |
| Primary | *Drosophila* | Anti-b-galactosidase rabbit IgG Polyclonal Antibody | 1:500 | Invitrogen | A11132 |
| Primary | *Drosophila* | Affinity purified Numb antibody (sheep | 1:1000 | Gifted by Jens Januschke | (Loyer et al., 2024) |
| Primary | Human | Anti-OSR2 antibody produced in rabbit | 1:250 | Fisher Scientific | PA5-106785 |
| Secondary | *Drosophila* & Human | Alexa Fluor 555 goat anti-rabbit IgG (H&L) | 1:500 | Invitrogen | A21428 |
| Secondary | *Drosophila* & Human | Alexa Fluor 488 goat-anti chicken IgG (H+L) | 1:500 | Invitrogen | A11039 |
| Secondary | *Drosophila* & Human | Alexa Fluor 488 goat-anti mouse IgG (H+L) | 1:500 | Invitrogen | A11001 |
| Secondary | *Drosophila* & Human | Alexa Fluor 546 goat-anti mouse IgG1 | 1:500 | Invitrogen | A21123 |
| Secondary | *Drosophila* & Human | Alexa Fluor 555 donkey-anti sheep IgG (H+L) | 1:500 | Fisher Scientific | A-21436 |
